# Nuclear Fetuin-A Drives Adipocyte Senescence via HIF-1α During Obesity

**DOI:** 10.1101/2025.08.18.670856

**Authors:** Debarun Patra, Ankit Vashisth, Soumyajit Roy, Palla Ramprasad, Shivam Sharma, Subrata Mishra, Biswa Mohan Prusty, Debasis Manna, Satpal Singh, Kulbhushan Tikoo, Suman Dasgupta, Durba Pal

## Abstract

Fetuin-A (FetA), a liver derived glycoprotein, has emerged from genome-wide association studies and epidemiological surveillance as a serum biomarker linked to obesity-driven type 2 diabetes mellitus (T2D), primarily due to its contribution to adipose tissue dysfunction. Here, we uncovered an eccentric role of nuclear FetA in visceral white adipocytes of obese T2D conditions. Hypoxia-inducible factor-1α (HIF-1α) facilitates the nuclear translocation of FetA via direct interaction, a process that promotes the emergence of a senescence-associated secretory phenotype (SASP). While nuclear co-localization of FetA and HIF-1α strongly promotes adipocyte senescence, silencing FetA alone is sufficient to prevent senescence, even in conditions of HIF-1α overexpression or lipid-rich hypoxic stress. Although nuclear FetA does not directly bind to DNA, it enhances HIF-1α transcriptional activity, potentiating the activation of senescence markers such as β-galactosidase and p53. Selective knockdown of FetA in obese mice notably reduced adipocyte senescence in visceral white adipose tissue (vWAT) and improved fasting glycemic control. Collectively, our findings reveal a previously unrecognized nuclear function for FetA in orchestrating adipocyte senescence in obesity, establishing nuclear FetA as a potential therapeutic target for obesity related metabolic diseases.

## Introduction

Obesity is a key aetiological factor for the hallmark of severe metabolic disorders such as insulin resistance and type 2 diabetes (T2D)(Kahn & Flier, 2000). In obese individuals, colossal expansion of adipose tissue (AT), mainly due to the hypertrophied adipocytes, coincided with the hypoxic, lipid-enriched adipose tissue microenvironment (AT*env*), which is causally linked to the remodelling and dysfunction of adipocytes(Patra *et al*, 2023). Previous reports indicated that elevated circulating levels of fetuin-A (FetA, also known as alpha-2 heremans schmid glycoprotein, AHSG), a member of the cystatin superfamily of cysteine protease inhibitors, are positively correlated with obesity progression and the development of obesity-related metabolic disorders including T2D(Kröger *et al*, 2018). Interestingly, epidemiological studies have been shown that the higher systemic levels of FetA are coupled with the incidence of T2D in aged individual irrespective of their sex or race(Ix *et al*, 2012). The FetA gene is mapped to the 3q27 region of human genome, a well-established T2D and metabolic syndrome susceptibility locus(Mathews *et al*, 2002a) and the ablation of FetA gene in mice confers resistant to weight gain on a high-fat diet along with the improvement of insulin sensitivity(Mathews *et al*, 2002b). FetA is mainly secreted from hepatocytes, however, its secretion can also be influenced by other cell types, including adipocytes, during obesity. It is associated with chronic low-grade inflammation in AT and insulin resistance by activating NF-κB-dependent signalling through direct interaction with toll-like receptor 4 (TLR4)(Pal *et al*, 2012). Additonally, FetA plays a direct role in promoting macrophages recruitment and polarization towards a proinflammatory state in obese AT(Chatterjee *et al*, 2013), and it contributes to insulin resistance by abrogating receptor tyrosine kinase activity^6^.

Senescence is a state of irreversible cell cycle arrest accompanied by cellular dysfunction, caused by different stressors such as telomere shortening, DNA damage, or oxidative stress, and is associated with the progression of aging and obesity(Tchkonia *et al*, 2010). Although senescent cells cease proliferation, they remain metabolically active and secrete various factors, including inflammatory cytokines and chemokines, characteristics of the senescence-associated secretory phenotype (SASP)(Wiley & Campisi, 2021; Van Deursen & M, 2014). Cellular senescence plays a crucial role in developing several metabolic disorders including non-alcoholic fatty liver disease, osteoporosis, cardiovascular diseases, and T2D(Palmer *et al*, 2019a). Moreover, clearance of senescent pancreatic β-cells has been shown to restore their function and improve blood glucose levels(Aguayo-Mazzucato *et al*, 2019). In obese individuals, particularly those with T2D, a substantial proportion of fully differentiated adipocytes manifest clear signs of cellular senescence and functional impairment. Recent literature indicates that obesity-driven hypertrophic expansion of adipocytes contributes to their dysfunction, accompanied by reduced insulin sensitivity and diminished lipid storage capacity(Palmer *et al*, 2019a). In addition, adipocyte senescence hampers the adipogenic potential and proliferation of resident progenitor cells(Nerstedt & Smith, 2023). Eliminating senescent cells or suppressing SASP expression may offer promising therapeutic strategies to restore or enhance adipocyte function, including improved insulin sensitivity.

In the present study, we discovered that FetA serves as a regulatory protein in adipocytes, with its nuclear migration facilitated by HIF-1α under the pathophysiological conditions of obese AT*env*, leading to the promotion of adipocyte senescence. Targeted inhibition of FetA effectively rescued adipocytes from SASP activation and the progression of cellular senescence. Thus, our study uncovers a novel mechanism of obesity-induced adipocyte senescence.

## Results

### Nuclear localization of FetA in adipocytes of obese visceral white adipose tissue

We have previously shown that obesity promotes the expression and secretion of hepatic FetA(Dasgupta *et al*, 2010; Chatterjee *et al*, 2013), which plays a crucial role in the induction of insulin resistance and T2D(Mori *et al*, 2006), however, its intracellular localization and regulatory role in adipocyte function remain unexplored. To address this issue, we examined the distribution of intracellular FetA in visceral white adipose tissue (vWAT) of Indian obese T2D patients (DM) compared to matched lean nondiabetic (ND) subjects. Immunofluorescence analyses of vWAT sections revealed an increased abundance of FetA in DM samples than in ND controls (Fig. 1a). Surprisingly, we noticed a marked increase in nuclear FetA in DM samples, as indicated by the higher co-localization of FetA and nuclear stain DAPI (Fig. 1b). Consistent with this, immunoblot analysis showed a significantly higher level of FetA in the nuclear fraction compared to cytosolic fraction of adipocytes isolated from DM patients (Fig. 1c), indicating the influence of obese AT*env* in nuclear occurrence of FetA protein. To apprehend whether hypoxic lipid-enriched (HL) microenvironment characteristics of obese adipose tissue^5^ could be accountable for the nuclear accumulation of FetA, we incubated differentiated 3T3-L1 adipocytes with 0.75mM palmitate under hypoxic conditions (1%O_2_). A significant enhancement of FetA protein levels was observed in both the cytosolic and nuclear fractions of HL-treated adipocytes, although the accumulation was more pronounced in the nuclear compartment (Fig. 1d). Immunofluorescence staining of adipocytes and isolated nuclei further demonstrated the nuclear enrichment of FetA following HL stimulation (Fig. 1e-h). However, pre-treatment of adipocytes with a nuclear transport inhibitor, ivermectin, markedly inhibited HL-induced nuclear accumulation of FetA (Fig. EV1A, B). Secretory FetA is a known glycoprotein that undergoes various post-translational glycosylation modifications(Pal *et al*, 2012; Lin *et al*, 2018a), and we detected glycosylation in the nuclear FetA, similar to its cytosolic counterpart (Fig. EV1C).

**Figure 1:**
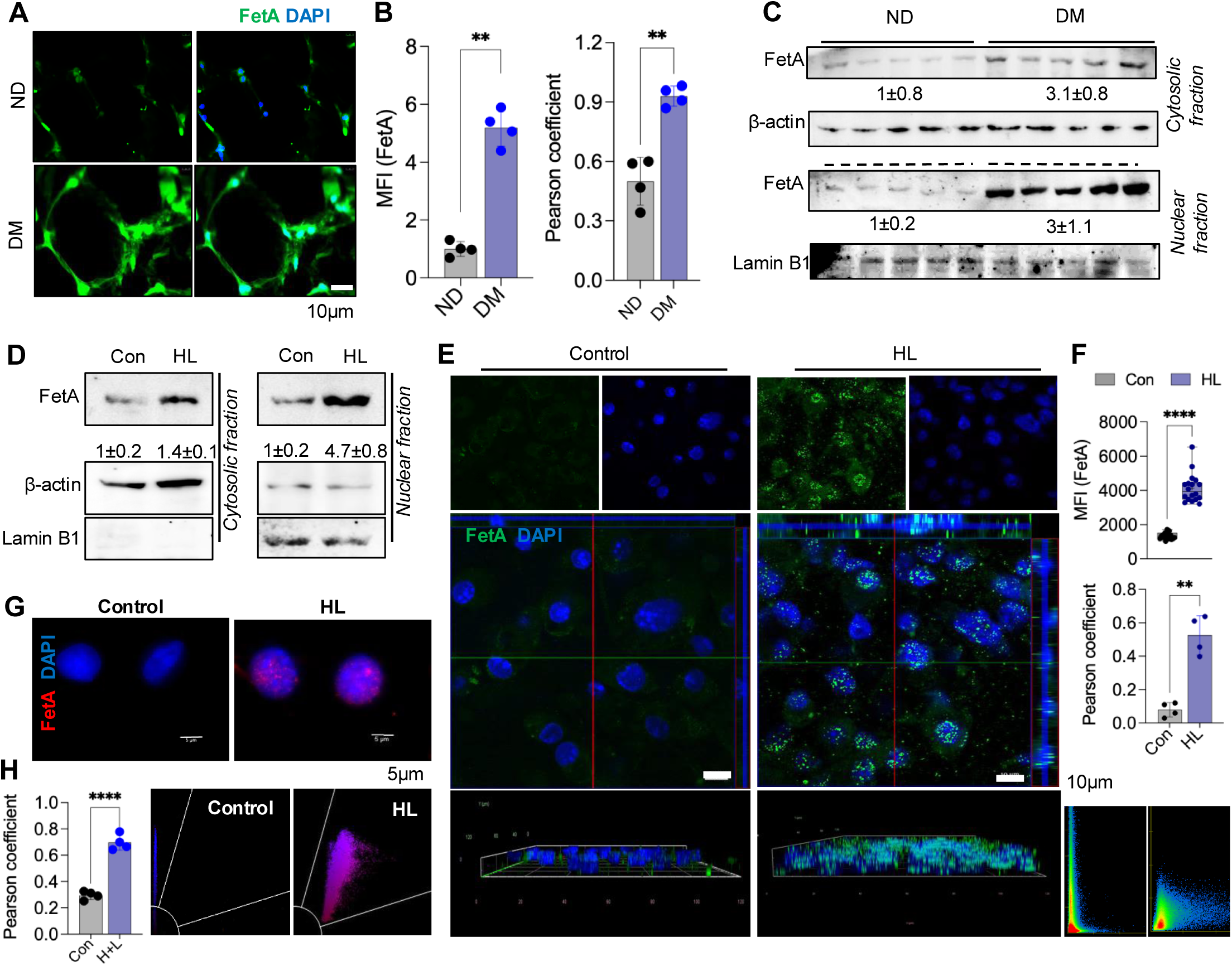
Nuclear localization of FetA in adipocytes from obese vWAT. A, Immunofluorescence analysis of FetA expression in histological sections of visceral white adipose tissue (vWAT) from lean non-diabetic (ND) and obese-Type 2 diabetic (OD) individuals. Scale bar, 10 μm. B, Quantification of FetA signal intensity and Pearson’s correlation coefficient from the images shown in (A). C, Immunoblot analysis of FetA protein levels in cytosolic and nuclear fractions of adipocytes isolated from vWAT of ND and OD individuals. D, Immunoblot analysis of FetA in cytosolic and nuclear fractions of 3T3-L1 adipocytes exposed to palmitate under hypoxic conditions (1% O₂, 16 hours; HL) or normoxia and no lipid. E, Confocal imaging of FetA localization in 3T3-L1 adipocytes under control or HL treatment. Scale bar, 10 μm. F, Quantification of mean fluorescence intensity (MFI) and Pearson’s correlation coefficient from the data shown in (E). G, Fluorescence imaging of isolated nuclei from control and HL-treated adipocytes showing FetA nuclear localization. Scale bar, 5 μm. H, Pearson’s correlation coefficient analysis of FetA nuclear localization from the data shown in (G).

All these results suggest that the pathophysiological obese AT*env* promotes the nuclear abundance of FetA in adipocytes.

### HIF-1**α** regulates the nuclear migration of FetA

Since FetA lacks any known putative nuclear localization signal (NLS) conferring for its ability to nuclear migration, we sought to investigate the mechanism of FetA nuclear translocation in adipocytes under the obese AT*env*. To explore the potential candidate responsible for the nuclear migration of FetA in adipocytes under obese AT*env*, the role of HIF-1α was examined, as we recently showed it’s activation in obesity potentiates adipose tissue inflammation and insulin resistance^2^. Moreover, recent reports also identify FetA as an evolutionarily conserved target gene of HIF-1α(Rudloff *et al*, 2021). We performed coimmunoprecipitation to examine the association of FetA with HIF-1α in AT*env*. In HL-exposed adipocytes, a significant enhancement of FetA and HIF-1α interaction was noticed, irrespective of whether immunoprecipitation was performed with anti-FetA or anti-HIF-1α antibodies, followed by immunoblotting with the corresponding counter-antibodies (Fig. 2A). Similar observations were made in isolated adipocytes from obese T2D patients and HFD mice (Fig. EV2A, B). To ensure that HIF-1α is critical for the nuclear translocation of FetA in obese adipocytes, 3T3-L1 adipocytes were transfected with HIF-1α siRNA and then incubated in the presence or absence of HL stimulation. Immunofluorescence staining and western blotting analyses corroborated the evidence that nuclear FetA was considerably reduced in HIF-1α-ablated adipocytes exposed to HL stimulation (Figs. 2B-D and Fig. EV2C). Further, the co-immunoprecipitation study demonstrated that the nuclear level of FetA and its association with HIF-1α were significantly abolished in the HIF-1α silenced adipocytes even in the presence of HL incubation (Fig. 2E and Fig. EV2D). However, forced expression of HIF-1α markedly increased the nuclear FetA level and its interaction with HIF-1α (Fig. 2F, G) in the absence of HL. These results suggest the obligatory requirement of HIF-1α in HL-induced nuclear migration of FetA. We then silenced FetA in HIF-1α-overexpressing cells to explore whether the localization pattern of HIF-1α is influenced by FetA. HIF-1α overexpression notably enhanced FetA protein levels, which is expected, as recent reports have shown that HIF-1α promotes FetA gene expression(Rudloff *et al*, 2021), however, FetA silencing was unable to restrict the nuclear migration of HIF-1α (Fig. 2H, I). These results indicate that HIF-1α nuclear translocation is independent of cellular FetA levels, although FetA migration to the nucleus is HIF-1α dependent.

**Figure 2:**
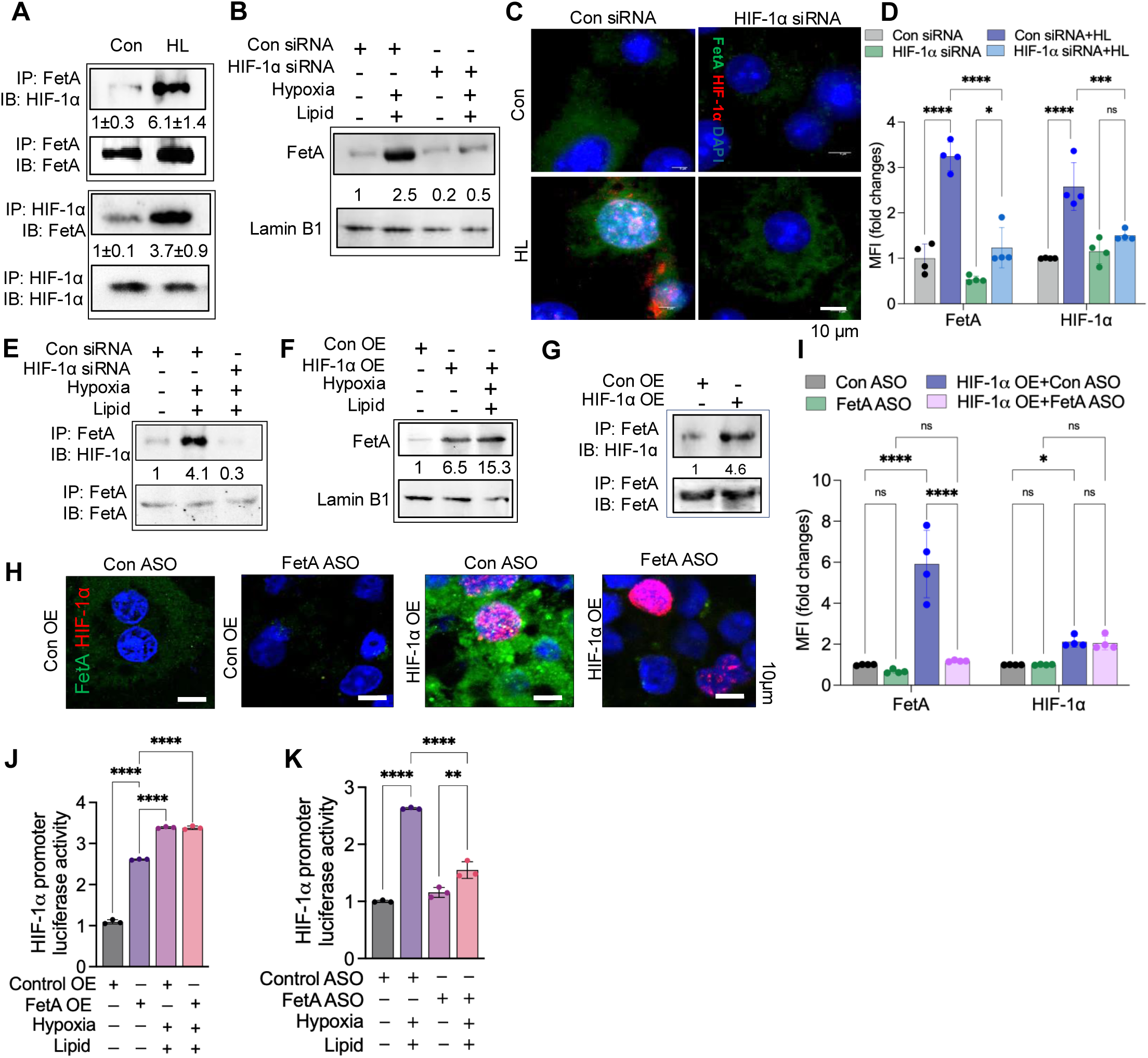
HIF-1α regulates the nuclear translocation of FetA. A, Immunoblot analysis of FetA and HIF-1α in total cell lysates and co-immunoprecipitation (co-IP) samples isolated from control and HL treated adipocytes using anti-FetA or anti-HIF-1α antibodies. B, Immunoblot analysis of FetA protein in control siRNA- and HIF-1α siRNA-transfected adipocytes exposed to HL or normoxic and no lipid conditions. C, Immunofluorescence imaging of FetA and HIF-1α in control siRNA- or HIF-1α siRNA-transfected adipocytes cultured with or without HL. Scale bar 10μm. D, Quantification of mean fluorescence intensity from images shown in (c). E, Immunoblot analysis of FetA and HIF-1α from nuclear co-IP samples isolated from control siRNA- or HIF-1α siRNA-transfected adipocytes treated with or without HL. F, Immunoblot analysis of FetA in the nuclear fraction of adipocytes transfected with control or HIF-1α overexpression (OE) plasmids under normoxic or HL conditions. G, Immunoblot analysis of FetA and HIF-1α in co-IP samples from nuclear lysates of control OE or HIF-1α OE plasmid-transfected adipocytes. H, Confocal imaging of FetA and HIF-1α in adipocytes transfected with HIF-1α OE plasmids or FetA antisense oligonucleotides (ASO), individually or in combination. Scale bar 10μm. I, Quantification of mean fluorescence intensity from the images shown in (H). J, HIF-1α luciferase reporter activity in FetA-overexpressing adipocytes under control or HL conditions. K, HIF-1α luciferase reporter activity in FetA-silenced adipocytes under control or HL conditions.

Chromatin immunoprecipitation (ChIP) assays revealed that nuclear FetA does not function as a transcription factor, as no direct interaction with genomic DNA, including the HIF-1α promoter region, was detected (data not shown). Interestingly, HIF-1α promoter-driven luciferase assays demonstrated that overexpression of FetA significantly enhanced HIF-1α promoter activity, independent of HL stimulation, whereas inhibition of FetA resulted in a marked decrease in promoter activity (Fig. 2J, K). These results suggest that nuclear FetA modulates HIF-1α activity indirectly, likely through the regulation of other intermediary proteins. Overall, our data reveal a critical role for adipocyte-derived nuclear FetA in modulating the hypoxic response in lipid-rich AT*env*.

### FetA directly binds with HIF-1α

To explore the physical interaction between FetA and HIF-1α, the pull-down assays and surface plasmon resonance (SPR) analyses were performed. For pull-down assay, HEK293 cells were co-transfected with HIF-1α-HA and FetA-HIS plasmids (Fig. EV3A), followed by immunoprecipitation of nuclear lysates with anti-HIS or anti-HA antibodies and subsequent probing with anti-HIF-1α or anti-FetA antibodies, respectively. Our data demonstrated a strong interaction between HIF-1α-HA and FetA-HIS proteins in the nuclear fraction upon their forced co-expression without any environmental stimuli (Fig. 3A). In SPR analyses, we first flowed varying concentrations of HIF-1α (analyte) over FetA (ligand) immobilized on a nitrilotriacetic acid (NTA) sensor chip surface. The SPR sensorgram displayed a concentration-dependent binding of HIF-1α to FetA with high binding affinity, as represented by the equilibrium dissociation constant (K_D_) value of 1.19×10^−8^M (Fig. 3B and Fig. EV3B). To confirm the interaction, increasing concentrations of FetA (analyte) were then passed over the HIF-1α (ligand) coated CM5 sensor chip surface. A similar pattern of concentration-dependent enhancement of high affinity binding between FetA and HIF-1α (K_D_ = 2.34×10^−8^ M) was observed (Fig. 3B and Fig. EV3B). These results suggest a strong physical interaction between FetA and HIF-1α proteins, which could be necessary for the nuclear localization of FetA. To investigate the specific molecular signatures of FetA and HIF-1α that could participate in their interaction, we collected the 3D structures of these proteins from the Alpha Fold Protein Structure Database (EMBL-EBI)(Jumper *et al*, 2021) and performed FetA-HIF-1α interaction analysis using the ClusPro web server(Desta *et al*, 2020). Since SPR analyses of full-length FetA (1-368 residues) and the C-terminal domain (CTD) of hypoxia-stable HIF-1α (579-826 residues) suggested a strong interaction between them, we therefore examined the specific residues of HIF-1α that could possibly interact with FetA through an *in-silico* bioinformatics study. Out of the several cluster models generated by our analysis, we deliberately selected the result file based on a thorough evaluation of balanced coefficients, taking into account the weighted score of -1946.6. Through bioinformatics study, we identified key amino acid residues of HIF-1α D743 and A746, among stretches of other residues (G735-S751, R781-Q794, and V802-N826) that are pivotal for its interaction with FetA (Fig. EV3C). Similarly, we found that S334, H335 and a region from G327-R340 of FetA appear critical for interaction with HIF-1α (Fig. EV3C). Our investigation unveiled certain critical amino acids that played a pivotal role in the intricate interaction between FetA and HIF-1α (Fig. 3C). To verify this, we performed deletion and point mutations on both FetA and HIF-1α plasmids at multiple sites (Fig. 3D, E). Intriguingly, we found that a point mutation in HIF-1α at 743D> L caused a sharp drop in its interaction with the WT FetA protein (Fig. 3F). Furthermore, a concentration gradient flow of mutant HIF-1α 743D>L (analyte) over wild-type (WT) FetA (ligand) immobilized on a NTA sensor chip surface showed a weaker binding affinity compared to WT HIF-1α, with at least an 8-fold decrease in the dissociation constant (K_D_ = 9×10^−8^ M) (Fig. 3G and fig. EV3D). On the other hand, we flowed mutant FetA proteins over a WT-HIF-1α-coated CM5 sensor chip surface at concentration of 45.45uM, and both point mutations, 334S>A and 335H>F, showed a sharp reduction in affinity compared to WT FetA (Fig. 3H), suggesting their critical role in stabilizing the interaction. We proceeded with the FetA 335H>F mutant, which showed the lowest response unit in the concentration gradient SPR analysis. The result revealed a 51.58-fold reduction in the dissociation constant (K_D_=120.7×10^−8^), suggesting a significantly weaker interaction affinity with the WT HIF-1α (Fig. 3I and fig. EV3D). Our results suggest that the D743 residue of HIF-1α and the H335 residue of FetA are critical for their interaction and play a pivotal role in facilitating nuclear migration of FetA.

**Figure 3:**
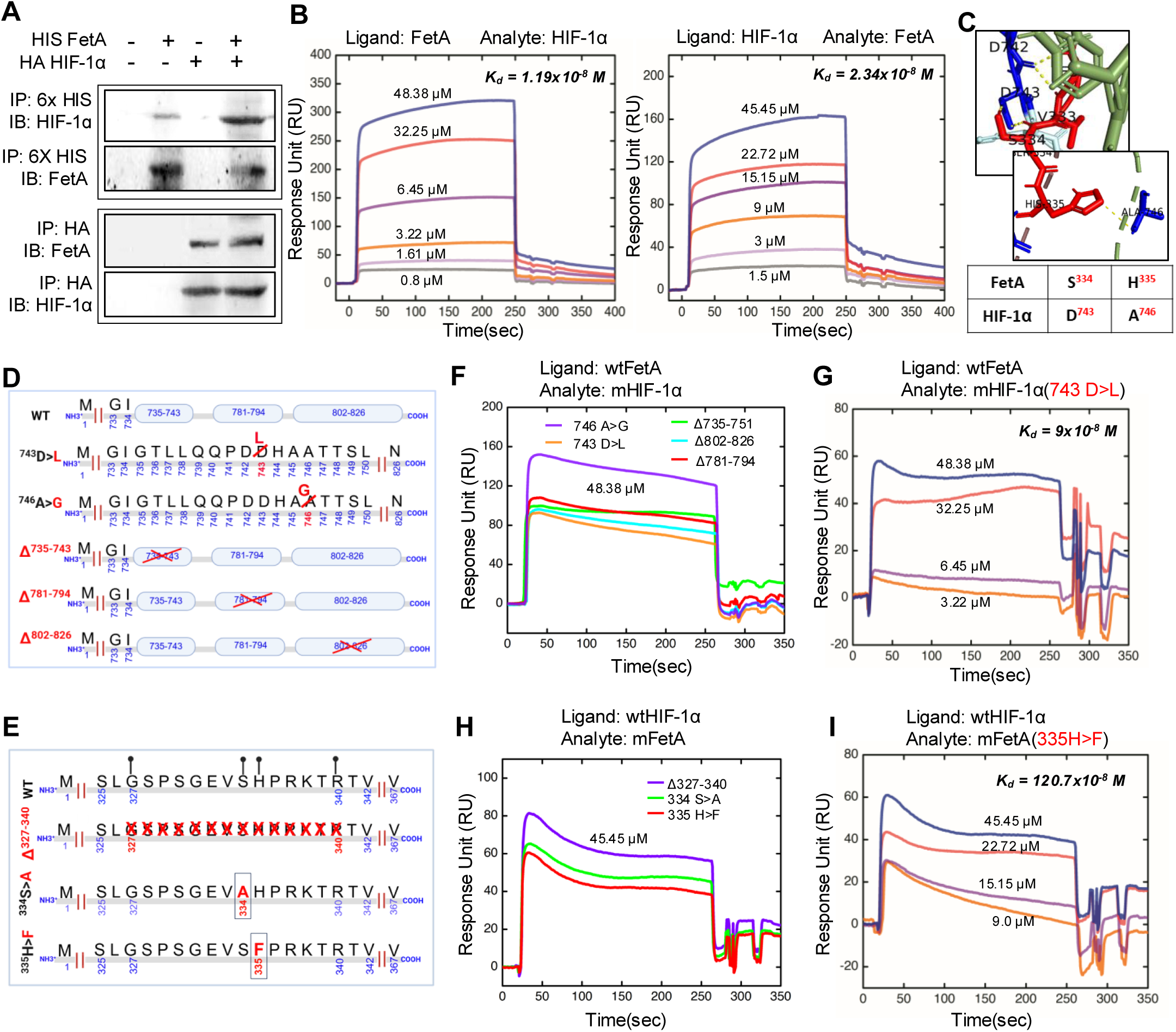
Fet-A directly interacts with HIF-1α. A, Co-immunoprecipitation of FetA and HIF-1α in HEK293T cells co-transfected with FetA-HIS and HIF-1α-HA expression constructs. Cell lysates were immunoprecipitated with anti-His or anti-HA antibodies, and HIF-1α and FetA expression was assessed by Western blotting. b, Surface plasmon resonance (SPR) sensorgrams showing real-time binding kinetics between FetA and HIF-1α. Either FetA (left) or HIF-1α (right) was immobilized on the sensor chip, and the indicated concentrations of the interaction partner were flowed over the surface. BSA was used as a negative control. Binding responses are shown in response units (RU) over time. C, Molecular model predicts active binding sites of human HIF-1α (blue) and human FetA (red). D, E, Schematic overview of the domain architecture of HIF-1α (D) and FetA (E), indicating the regions targeted for deletion or point mutation to map the interaction interface. F, H, SPR sensorgrams illustrating the effect of site-specific mutations on FetA–HIF-1α binding. Mutated HIF-1α proteins were flowed over immobilized WT-FetA at a concentration of 48.38 μM (F), and conversely, mutated FetA proteins were flowed over immobilized WT-HIF-1α at 45.45 μM (H). G, SPR sensorgrams showing binding responses of the HIF-1α 743 D>L point mutant protein titrated over immobilized WT-FetA. I, SPR sensorgrams showing binding responses of the FetA 335 H>F point mutant protein titrated over immobilized WT-HIF-1α.

### Nuclear FetA regulates HIF-1α and induces adipocyte senescence

The intriguing discovery of FetA nuclear translocation in adipocytes during obesity prompted us to investigate its functional relevance, particularly in the context of adipocyte senescence, a process increasingly recognized as a contributor to metabolic dysfunction in obesity and aging(Tchkonia *et al*, 2010; Palmer *et al*, 2019b). To assess whether nuclear FetA plays a role in the induction of senescence induction, we first evaluated senescence-associated β-galactosidase (SA-β-gal) activity in adipocytes exposed to a lipid-rich, hypoxic (HL) AT*env*. HL treatment significantly elevated SA-β-gal activity compared to controls, consistent with the induction of cellular senescence (Fig. EV4A).

Supporting this observation, RT-qPCR analyses revealed a robust upregulation of SASP markers, including β-gal, PAI-1, p53, p21, TNF-α, MMP3, CCL20, IL-8, FAS, TIMP2, and CCND1 in HL-treated adipocytes (Fig. EV4B). Confocal imaging further confirmed that HL-induced senescence was accompanied by enhanced nuclear accumulation of FetA alongside elevated β-gal expression (Fig. EV4C, D).

To dissect the mechanistic role of FetA, we performed gain-of-function experiments by overexpressing FetA and HIF-1α, individually or in combination in adipocytes. Co-overexpression of FetA and HIF-1α markedly increased nuclear FetA levels and strongly induced the expression of SASP markers β-gal, TNF-α, IL-6, and MMP3 (Fig. 4A, B). Notably, FetA overexpression alone did not induce senescence, highlighting the requirement for HIF-1α-mediated nuclear translocation of FetA in this process (Fig. EV4E). Consistently, SA-β-gal staining intensity was substantially higher in adipocytes co-expressing FetA and HIF-1α compared to controls (Fig. 4C).

**Figure 4:**
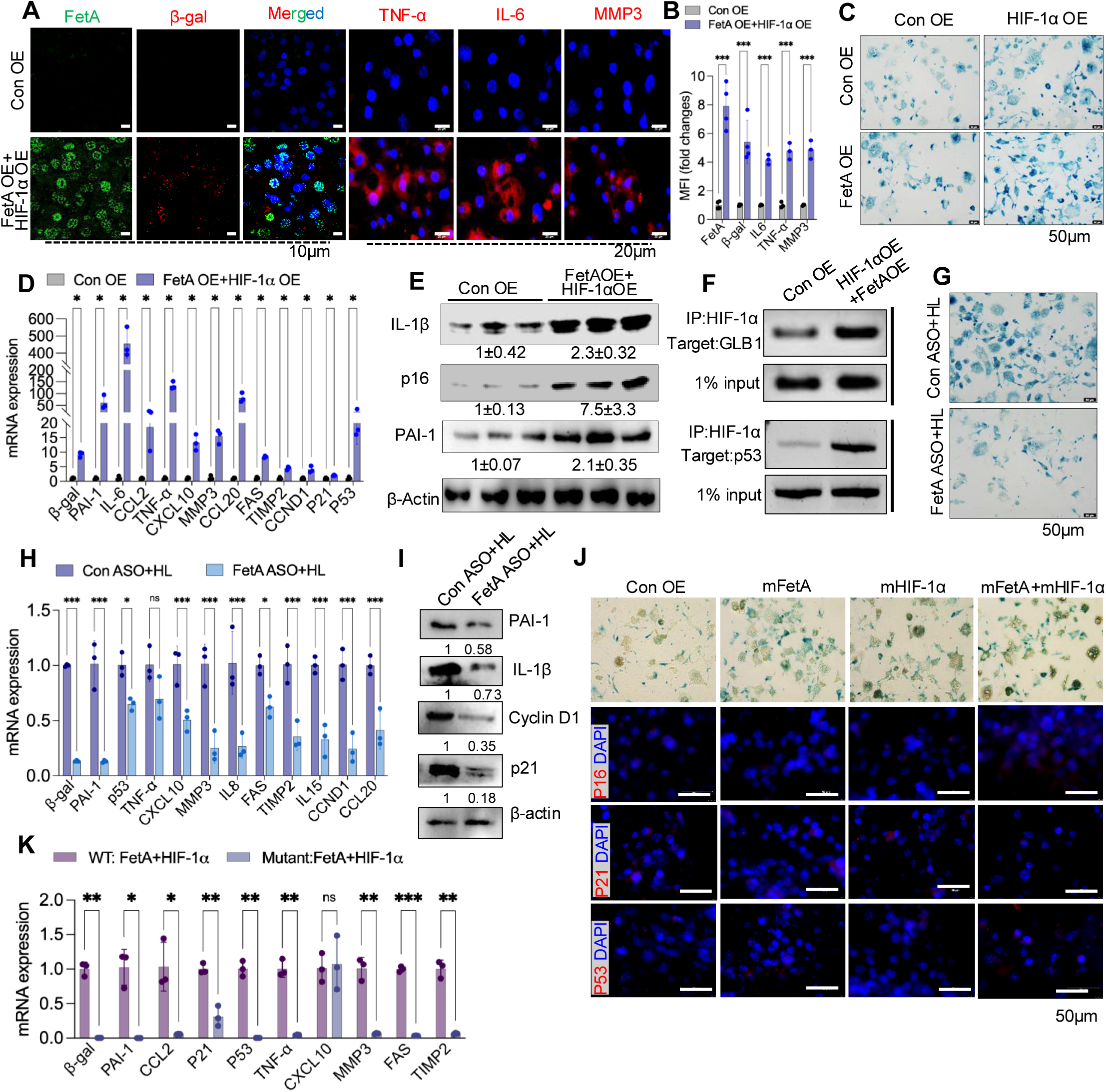
Nuclear FetA induces adipocyte senescence. A, Confocal imaging of senescence-associated secretory phenotype (SASP) markers β-gal, TNF-α, IL-6, and MMP3 (red) along with FetA (green) in adipocytes transfected with control overexpression or HIF-1α and FetA overexpression (OE) plasmid in combination. Scale bar, 10 μm & 20 μm. B, Quantification of mean fluorescence intensity corresponding to data shown in (A). C, Senescence-associated β-galactosidase (SA-β-gal) staining and imaging of adipocytes transfected with HIF-1α or FetA OE plasmid, either alone or together. Scale bar, 50 μm. D, qRT-PCR analysis of SASP gene expression in adipocytes co-transfected with HIF-1α and FetA OE plasmids compared to control. E, Immunoblotting analysis and densitometric quantification of SASP markers in HIF-1α and FetA co-transfected adipocytes versus control. F, Chromatin immunoprecipitation (ChIP) analysis for GLB1 (β-gal) and p53 promoter occupancy by HIF-1α in adipocytes co-transfected with or without HIF-1α OE and FetA OE plasmids. G, SA-β-gal staining in control and FetA-silenced adipocytes treated under lipid-rich hypoxic conditions (HL). Blue precipitate indicates senescent cells. Scale bar, 50 μm. H, mRNA expression analysis of SASP markers in control versus FetA-silenced adipocytes exposed to HL. I, Immunoblotting and densitometric quantification of SASP markers in control and FetA-silenced adipocytes under HL treatment. J, SA-β-gal staining and immunofluorescence analysis of SASP markers in adipocytes transfected with mutated HIF-1α and/or mutated FetA constructs. Scale bar, 50 μm. K, RT-qPCR analysis of SASP genes expression in adipocytes co-transfected with mutated FetA and mutated HIF-1α plasmids.

Further, co-transfection of FetA and HIF-1α plasmids amplified the expression of SASP genes and proteins, as confirmed by RT-qPCR and immunoblotting (Fig. 4D, E). ChIP assays revealed enriched HIF-1α binding to the promoters of SASP genes (β-gal, p53) under these conditions, suggesting a direct transcriptional role for the FetA-HIF-1α complex (Fig. 4F).

*Loss-of-function* studies using FetA-specific antisense oligonucleotides (ASO) markedly reduced HL-induced SA-β-gal activity and suppressed both SASP gene and protein expression (Fig. 4G-I and Fig. EV4F, G). Moreover, silencing either FetA or HIF-1α abolished SA-β-gal induction, even upon overexpression of their binding partner (Fig. EV4H, I). Finally, mutational disruption of the FetA (335H>F)-HIF-1α (743 D>L) interaction, identified via SPR analysis, also abrogated the senescence phenotype, reinforcing the quintessential role of this protein complex in the regulation of adipocyte senescence (Fig. 4J, K).

Collectively, these results highlight a key regulatory axis involving HIF-1α-driven nuclear translocation of FetA, which cooperatively induces senescence in adipocytes exposed to the AT*env*. Targeted disruption of this pathway may provide new therapeutic opportunities to mitigate obesity-associated AT dysfunction.

### FetA inhibition protects against obesity-induced adipocyte senescence

To investigate the in vivo relevance of nuclear FetA in adipocyte senescence, we established a diet-induced obesity model by subjecting mice to HFD for 16 weeks, followed by treatment with either control morpholino or FetA-targeting morpholino over a 14-day period (Fig. 5A). Consistent with previous reports(Wan *et al*, 2024; Smith *et al*, 2021; Li *et al*, 2021; Spinelli *et al*, 2023), histological analysis of vWAT from obese T2D patients and HFD-fed mice revealed a pronounced increase in SASPs (Fig. EV5A–D). Silencing of FetA in the vWAT of obese mice was confirmed at both mRNA and protein levels following FetA morpholino administration (Fig. 5B and Fig. EV5E, F). Notably, suppression of FetA led to a marked decrease in the burden of senescent cells, as evidenced by reduced SA-β-gal staining and diminished immunohistochemical signals for senescence markers (β-gal, PAI-1, p16, Cyclin D1) in vWAT sections (Fig. 5C-F). FetA silencing also led to a substantial reduction in fasting blood glucose levels, without affecting body weight or altering the gross tissue architecture of vWAT (Fig. 5G, H and Fig. EV5G). Further molecular analyses demonstrated a significant decrease in the SASP-related genes (β-gal, p53, CCND1, p21, MMP3, FAS, TIMP2, IL6) and proteins (TNF-α and IL-1β) in adipocytes isolated from the vWAT of FetA-silenced mice (Fig. 5I, J). Collectively, these findings uncover a previously unrecognized role for nuclear FetA as a key critical regulator of adipocyte senescence in obesity-associated metabolic dysfunction.

**Figure 5:**
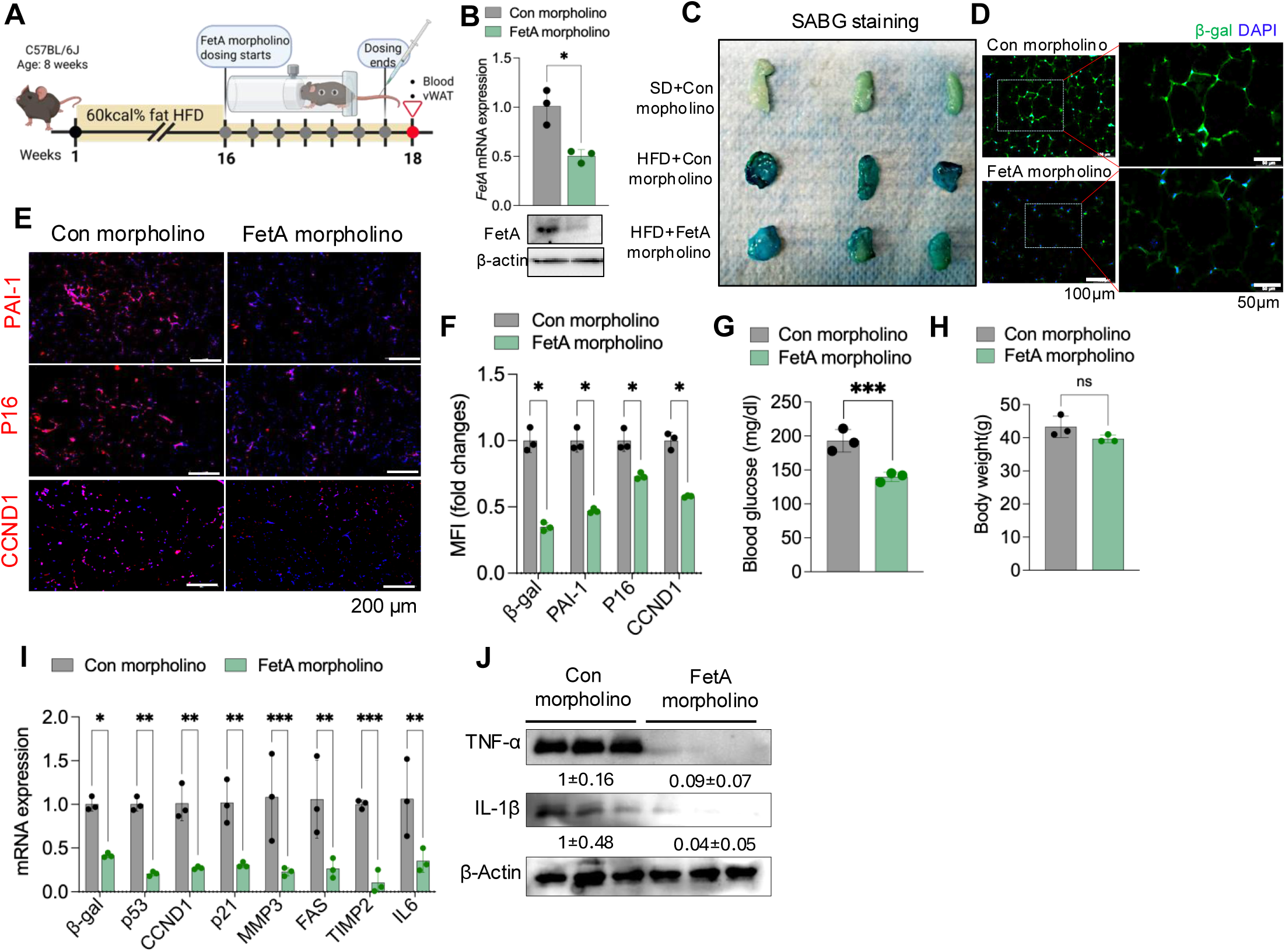
FetA inhibition protects against obesity-induced adipocyte senescence. A, Schematic representation of the experimental design depicting the development of diet-induced obesity by HFD feeding and subsequent treatment with control or FetA targeting morpholinos. B, Quantification of FetA mRNA and protein levels in vWAT by qRT-PCR and immunoblotting, respectively. C, Representative images of SA-β-gal staining in vWAT from standard diet (SD) and HFD mice treated with control or FetA morpholino (n=3). D, E, Representative immunofluorescence images of β-gal, PAI-1, p16, and Cyclin D1 in vWAT sections from control or FetA morpholino-treated HFD mice. F, Quantification of mean fluorescence intensity corresponding to the data shown in (D, E). G, Fasting blood glucose levels in HFD mice treated with control or FetA morpholino. H, Body weight measurements. I, qRT–PCR analysis of SASP markers (β-gal, p53, CCND1, p21, MMP3, FAS, TIMP2, IL-6) in adipocytes isolated from vWAT of HFD mice treated with control or FetA morpholino. J, Immunoblot analysis and quantification of TNF-α and IL-1β in vWAT adipocytes from control or FetA morpholino-treated HFD mice.

## Discussion

Obesity is well known to elevate the expression and secretion of FetA from hepatocytes and adipocytes, contributing significantly to the development of insulin resistance and T2D (Trepanowski *et al*, 2015). While previous studies including our own have primarily focused on the extracellular role of FetA, we previously demonstrated that FetA levels are elevated in the serum of obese individuals due to increased hepatic secretion(Pal *et al*, 2012; Trepanowski *et al*, 2015; Bourebaba & Marycz, 2019). This circulating FetA contributes to chronic inflammation by directly binding to TLR4 and activating the NF-κB signaling pathway. Importantly, inhibition of FetA in obese mouse models was shown to significantly reduce systemic inflammation and improve insulin sensitivity. In the current study, we extend these findings by identifying a novel intracellular role of FetA specifically its nuclear localization in adipocytes which appears to be critically involved in driving adipocyte senescence, a hallmark of dysfunctional adipose tissue in obesity. Our findings address a more comprehensive understanding of FetA’s pathogenic function.

We identified HIF-1α as the upstream regulators potentially mediating the nuclear translocation of FetA under the influence of the obese adipose tissue environment (AT*env*), we investigated the involvement of HIF-1α. This was based on our earlier work demonstrating that HIF-1α activation contributes to adipose tissue inflammation and insulin resistance in obesity(Takikawa *et al*, 2016; Lee *et al*). HIF-1α belongs to a family that includes the oxygen-sensitive HIF-1α, -2α, and -3α which form heterodimers with a constitutively expressed non-oxygen responsive HIF-1β subunit of the PAS family (PER, AHR, ARNT, and SIM)(Weidemann & Johnson, 2008). Under normoxic conditions, HIF-1α is rapidly degraded by prolyl hydroxylase domain (PHD) containing enzymes which hydroxylate proline residues present in its oxygen-dependent death domain. This hydroxylation facilitates recognition and binding by the E3 ubiquitin ligase complex, the von Hippel–Lindau tumor suppressor protein, leading to HIF-1α degradation. During hypoxic conditions, hydroxylation of HIF-1α is inhibited, resulting in its accumulation and nuclear localization followed by dimerization with HIF-1β. The HIF-1α/HIF-1β heterodimer binds to the core pentanucleotide sequence (RCGTG) present in the hypoxia response elements (HREs) of target genes and governs their expression(Weidemann & Johnson, 2008; Majmundar *et al*, 2010; Greer *et al*, 2012).

The sequence analysis of HIF-1α revealed that S576 and S657 are known to contribute to its stabilization, whereas the K719T mutation markedly reduces its nuclear migration. Additionally, mutations at the conserved ‘LPXL sequence motif’ at positions 792-795, particularly L795, as well as at C800 and N803 are detrimental to its transactivation potential mediated by the interaction with different cofactors including p300/CBP(Ema *et al*, 1999; Freedman *et al*, 2002; Xu *et al*, 2010; Lando *et al*, 2002). Our integrative approach combining in-silico predictions and co-immunoprecipitation assays demonstrated a direct physical interaction between FetA and HIF-1α within the obese AT*env*. Mutation analyses performed in-silico targeting key residues on FetA (S334A and H335F) and HIF-1α (D743L and A746G) highlighted their critical roles in mediating this interaction with their respective wild-type counterparts (WT-HIF-1α and WT-FetA), as shown in Fig. EV3E. These findings showed the biophysical interaction between FetA and HIF-1α in white adipocytes. The interaction was further validated through SPR analysis, which revealed a strong binding affinity between these two proteins.

FetA undergoes post-translational modifications in the endoplasmic reticulum and golgi apparatus, beginning with N-glycosylation and O-glycosylation, respectively together contributing to more than 23% of the protein’s carbohydrate content(Lin *et al*, 2018b). Our previous work demonstrated that terminal β-glycosides on secreted FetA are essential for its interaction with TLR4(Pal *et al*, 2012). However, the glycosylation status of the nuclear-FetA remains unknown. Our PAS staining analysis on nuclear and cytosolic FetA did not reveal any significant changes in glycosylation patterns. These findings underscore the need for more detailed investigations into the total carbohydrate composition of nuclear, cytosolic, and secretory forms of FetA, particularly in the context of obesity-associated adipocyte dysfunction.

While the AT*env* are known to promote inflammation, macrophage polarization, and insulin resistance(Patra *et al*, 2023), the physiological relevance of the FetA–HIF-1α interaction and their nuclear co-localization remained unclear. We observed HL treatment led to heightened expression of SASP markers, which strongly correlated with increased nuclear localization of FetA in adipocytes. Functionally, silencing or blocking nuclear translocation of FetA in mature adipocytes resulted in reduced SA-β-gal activity under obese conditions. Furthermore, in vivo administration of FetA morpholino in HFD mice significantly decreased SASP genes expression, underscoring the previously unrecognized pathological relevance of nuclear FetA in driving adipocyte senescence.

These findings strongly suggest a remarkable role for nuclear FetA in driving obesity-induced adipocyte dysfunction. However, to solidify these mechanisms, studies using FetA knockout models will be essential. Our data also indicates that the FetA-HIF1α interaction contributes to the induction of the SASP and promotes adipocyte senescence. This highlights the need for further investigation, particularly into therapeutic approaches that could selectively block FetA’s nuclear translocation. Additionally, an important question for future investigation is whether FetA also influences hepatic and subcutaneous white adipose tissue, potentially broadening our understanding of its systemic role in metabolic disease.

In summary, our study is the first to report the nuclear presence of FetA in adipocytes, revealing that its translocation is facilitated by direct interaction with HIF-1α. This nuclear interaction appears to be a pivotal driver of the adipocyte senescence program during obesity, offering new mechanistic insights and potential therapeutic targets for metabolic diseases associated with obesity.

## Acknowledgments

We thank NIPER SAS Nagar, Animal House Facility for mice experiments and IIT Ropar for providing instrumentation facilities. This study was supported by the SERB-Early Career Research Grant (Project No.: ECR/2017/000892), Govt. of India, DBT-Wellcome Trust Intermediate Fellowship to Du.P. and the DBT-Twining project (Project No.:BT/PR24700/NER/95/819/2017) to S.D. and Du.P. De.P., and P.R., acknowledge the IIT Ropar and the Ministry of Education, Govt. of India for their research fellowships. A.V., acknowledge the CSIR-HRDG, Govt. of India for research fellowship.

## Authorship contribution statement

Conceptualization: De.P., S.D., Du.P.; Methodology: De.P., S.D., Du.P.; Investigation: De.P., A.V., P.R., S.R., Sh.S., S.M., B.M.P., Sa.S., S.D. ; Validation: De.P., A.V., P.R., S.R., Sh.S., S.M., B.M.P., D.M., Formal Analysis: De.P., A.V., S.D., Du.P.; Writing–Original Draft: De.P., S.D., Du.P.; Writing–Review and Editing: De.P., K.T., S.D., Du.P.; Funding Acquisition: Du.P., S.D.; Resources: K.T., S.D., D.M., Du.P.; Supervision: S.D., Du.P.

## Data, Materials, and Software Availability Statement

All study data are included in the article and/or *Expanded View Figure and Table*. Du.P. is the guarantor of this work and, as such, has full access to all the data in the study and takes responsibility for the integrity of the data and the accuracy of the data analysis.

## Materials and Methods

### Materials

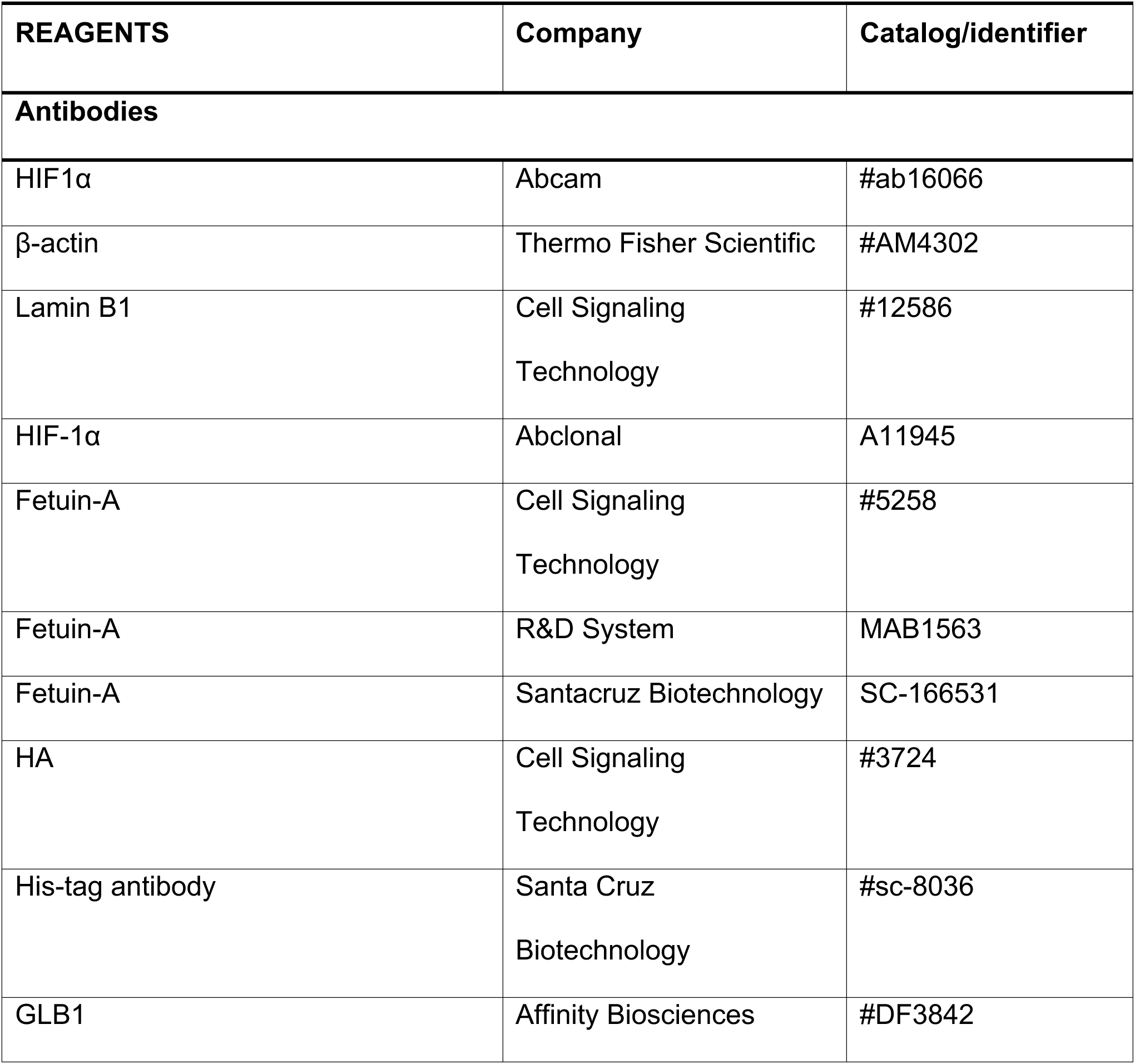
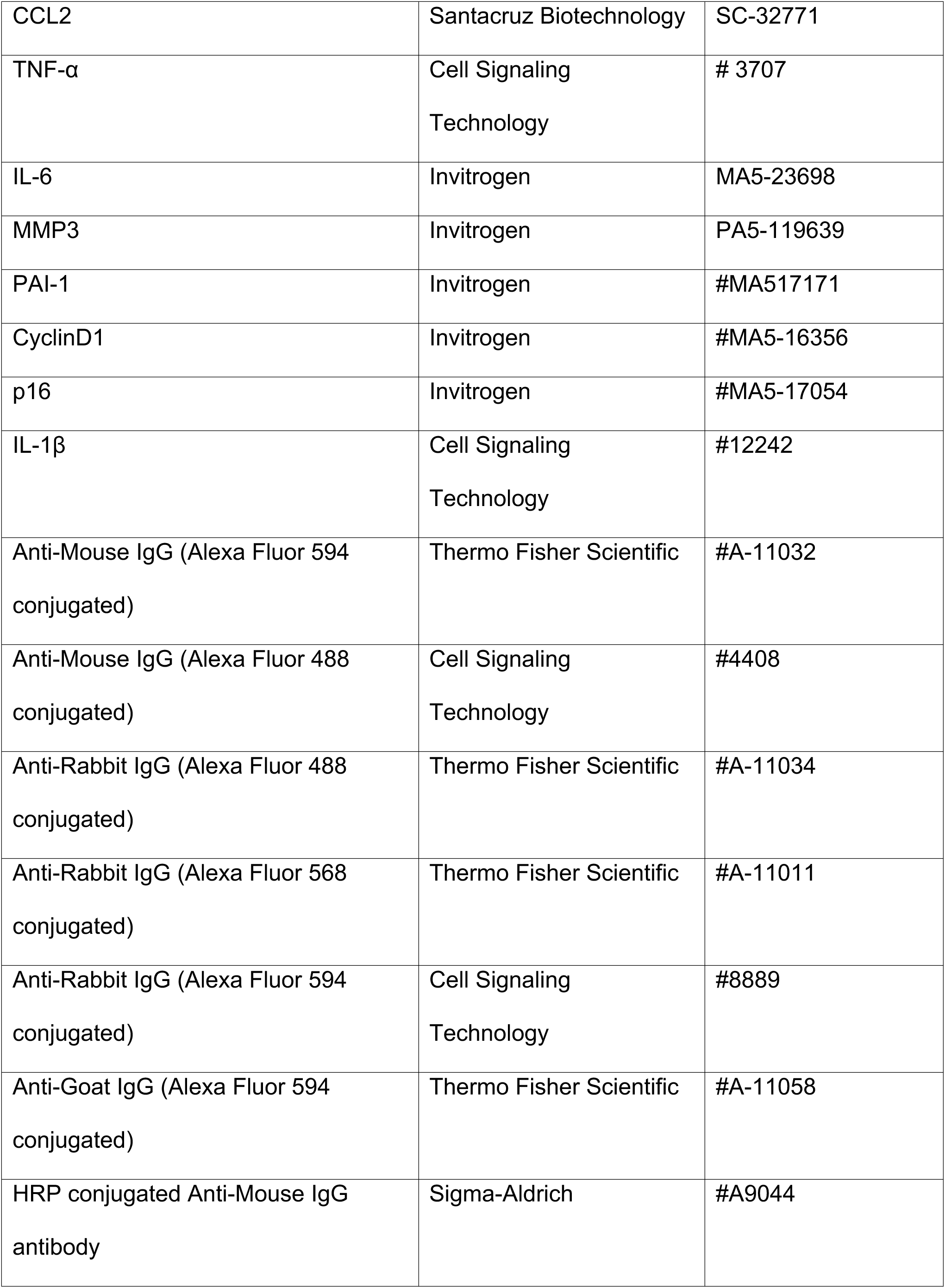
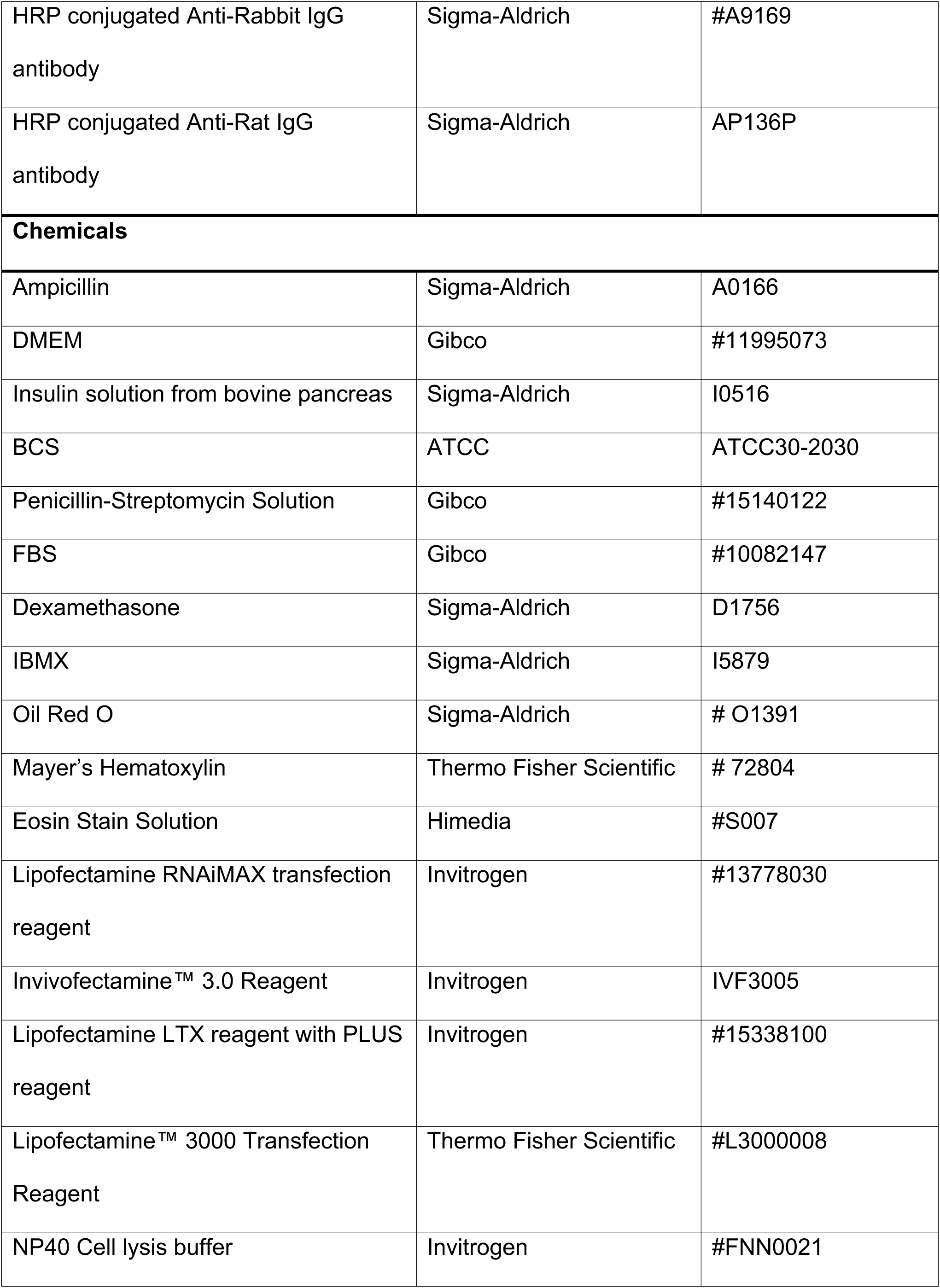
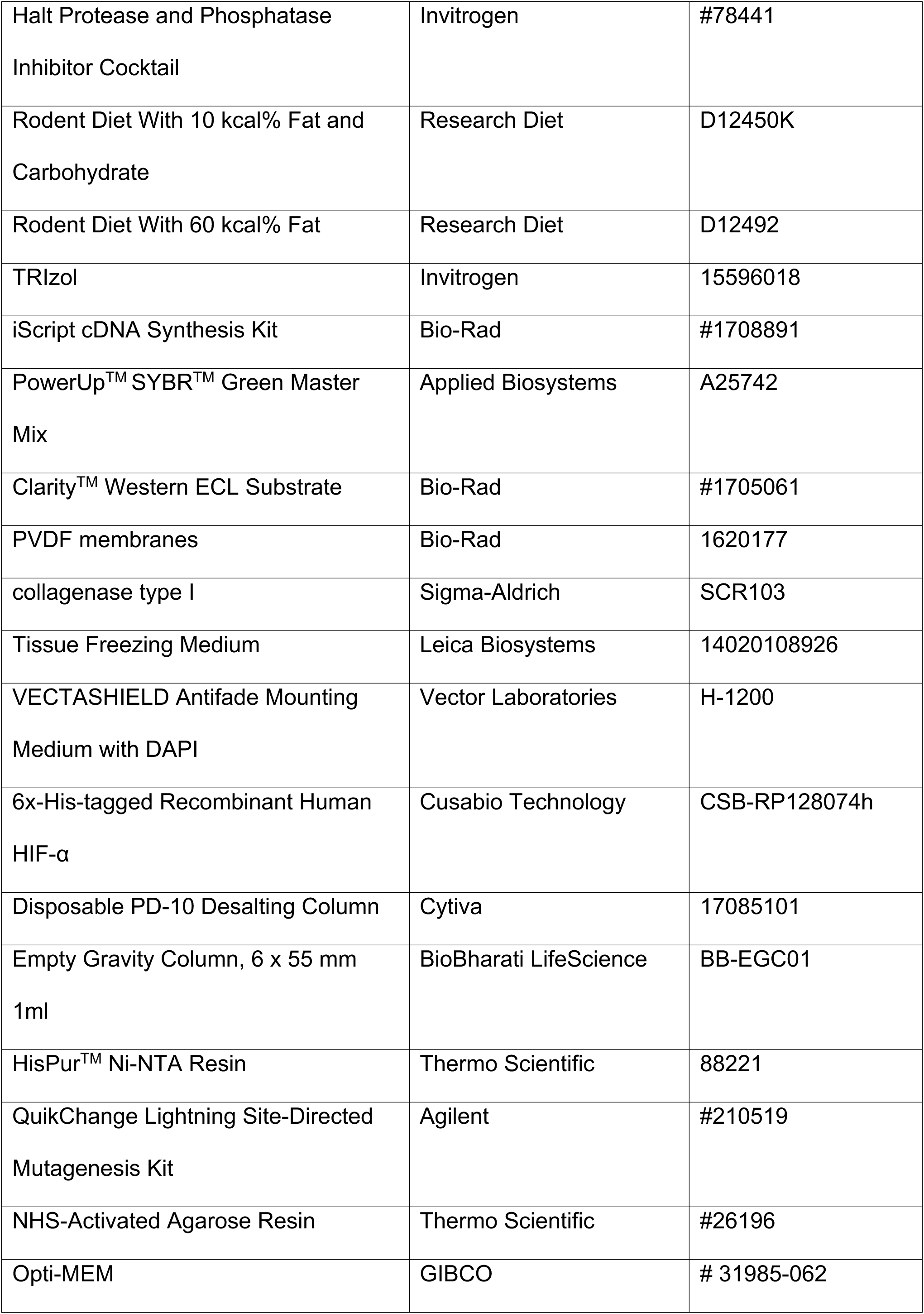
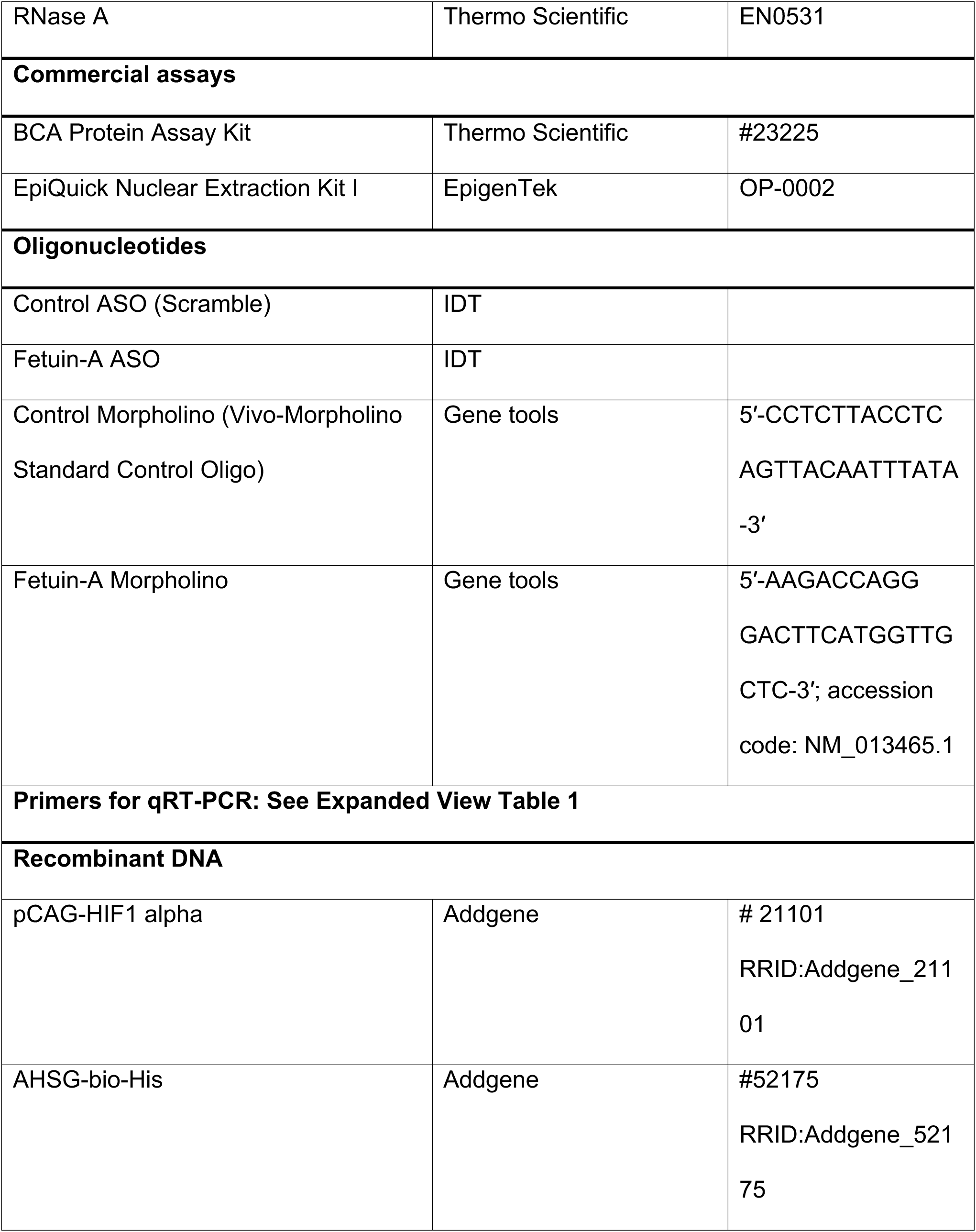
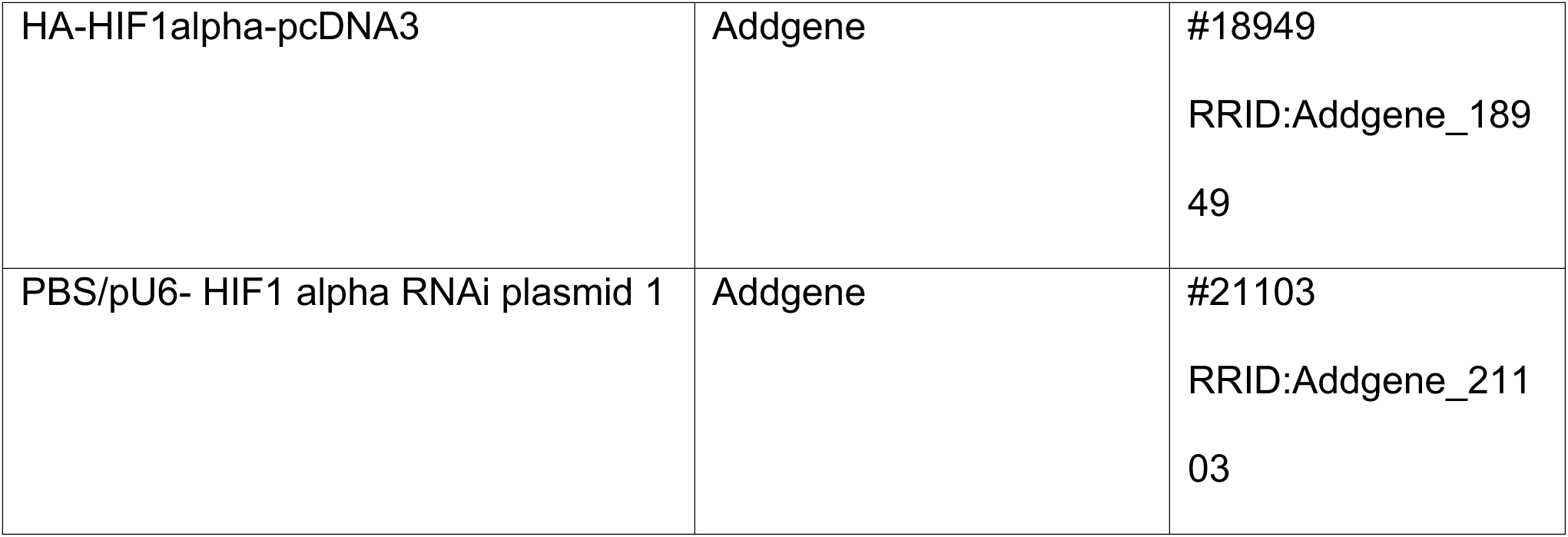

### Methods

#### Cell line

3T3-L1 cells were cultured in a growth medium containing DMEM supplemented with 10% BCS and 1% Penicillin-Streptomycin Solution. 3T3-L1 preadipocytes were differentiated into adipocytes following chemically induced differentiation protocol of ATCC. Once full confluency was reached, differentiation medium containing DMEM with high glucose, HI-FBS, 1.0 µM Dexamethasone, 0.5 mM IBMX, and 1µg/mL Insulin was added. After induction, differentiation medium was replaced by adipocyte maintenance medium containing DMEM with high glucose, 10% HI-FBS, and 1 µg/mL Insulin. We incubated adipocytes with a fixed concentration of palmitate (0.75mM) and exposed them to hypoxia condition (1% O_2_, and 5% CO_2_) for different time periods in the Heracell^TM^ VIOS 160i incubator. Cell culture reagents details were presented in materials Table.

#### Animals

Wild-type C57BL/6J male mice aged 4-5 weeks and weighed 18-22 g were kept in the NIPER Mohali animal house facility for 5-6 days in 12 light/dark cycle at 23 ± 2°C with relative humidity 55 ± 5% and fed with normal pellet diet and water ad libitum. For the development of a diet-induced obese and insulin resistance model, C57BL/6J mice were fed with high-fat diet (HFD) pellets having 60% kcal of fat for 12 weeks. All other mice were fed with standard diet (SD) having 10% kcal of fat for 12 weeks. All experimental animals had free access to sterilized water and food. The blood glucose levels were measured regularly with Accu-Chek glucometer (Roche). Mice fed with HFD diet for 12 weeks were considered for fetuin-A silencing. For this, Control morpholino or Fetuin-A morpholino (25nM per animal) was administered via the tail-vein of HFD mice, 7 times in 14 days. On day 14, animals were utilized for different experiments. All animal experiments were performed following the guidelines prescribed by and with the approval of the institutional animal ethics committee (IAEC) of the NIPER S.A.S. Nagar, Punjab.

#### Human participants

The study population was categorized based on BMI and blood glucose level. Study subjects had BMI 18–25 kg m^-2^ and fasting blood glucose level <85 mg dl^-2^ were considered as lean non-diabetic group (ND, n=5), whereas, patients with BMI >30 kg m^-2^ and fasting blood glucose level > 120 mg dl^-2^ were considered as obese diabetic group (DM, n=5) as presented in the Expanded View Table 3. In this study, surgically dissected visceral white adipose tissue (vWAT) samples and blood samples were collected from the patients who underwent abdominal surgery at the Dayanand Medical College & Hospital, Ludhiana, Punjab. The study protocol for the use of human blood and tissue samples was approved by the Institute Ethics Committee (IEC), Dayanand Medical College & Hospital, Ludhiana, Punjab. The written informed consent from all participants was obtained in this study. Data presented in Expanded View Table 2.

#### Plasmids and ASOs transfection

For the transfection of siRNA or overexpression plasmid, Lipofectamine LTX/Plus Reagent was used according to the manufacturer’s protocol. Briefly, cells were cultured in a 6-well plate in an antibiotic-free complete growth medium before transfection. For each well, 5μg of siRNA or overexpression plasmid and Lipofectamine LTX/Plus Reagent were added separately into the Opti-MEM serum free medium. Both these solutions were mixed and incubated for 5 min. The transfection mixture was then added to the cells containing complete growth medium and incubated for 48 h. Cells were washed, fresh media was added and used for different treatments.

pCAG-HIF1 alpha was a gift from Connie Cepko (Addgene plasmid # 21101 ; http://n2t.net/addgene:21101 ; RRID:Addgene_21101)(Chen & Cepko, 2009), AHSG-bio-His was a gift from Gavin Wright (Addgene plasmid # 52175 ; http://n2t.net/addgene:52175 ; RRID:Addgene_52175)(Sun et al, 2015), (Addgene plasmid # 18949 ; http://n2t.net/addgene:18949 ; RRID:Addgene_18949)(Kondo et al, 2002), PBS/pU6-HIF1 alpha RNAi plasmid 1 was a gift from Connie Cepko (Addgene plasmid # 21103 ; http://n2t.net/addgene:21103 ; RRID:Addgene_21103)(Chen & Cepko, 2009).

Control ASO or FetA ASO was transfected using Lipofectamine^TM^ RNAiMAX transfection reagent was used according to the manufacturer’s protocol. Briefly, 3T3-L1 adipocytes(differentiated) were cultured in a 6-well plate in an antibiotic-free complete growth medium before transfection. For each well, 50 nM control ASO or 100 nM of FetA ASO in Lipofectamine RNAi MAX reagent was added separately into the OptiMEM serum-free medium. Both these solutions were mixed and incubated for 5 min. The transfection mixture was added to the cells containing complete growth medium and incubated for 48 h. After 48 h of transfection, cells were washed; a fresh complete growth medium was added and used for different treatments.

#### Adipocyte nuclei staining and nuclear protein isolation

Control and treated adipocytes were washed twice with DPBS and incubated with chilled ethanol for 5 minutes, followed by the washing with DPBS, and then added with chilled hypotonic lysis solution (20 mM HEPES pH 7.5 containing 5 mM Sodium chloride, 0.4% NP-40, along with a protease and phosphatase inhibitors cocktail). Cell lysates were harvested and passed through the 26-gauge needle for few times and then centrifuged at 600 x g for 20 minutes at 4°C to pellet down the nuclei. The supernatant containing cytoplasmic lysate was removed and the nuclei pellet was washed 2-3 times with the hypotonic lysis solution resuspended and incubated first with 0.2% triton X-100 containing PBS for 5 minutes, then with 3% BSA solution for 15 minutes. The nuclei suspension was probed with anti-FetA primary antibody followed by the fluorochrome-tagged secondary antibody. The stained nuclei were counterstained with 100 nM DAPI solution and subjected to fluorescent imaging. The adipocytes were also subjected to nuclear protein isolation using the EpiQuick Nuclear Extraction Kit following manufacturer protocol.

#### Co-immunoprecipitation

The co-immunoprecipitation study was performed on the adipocytes isolated from vWAT of humans, mice, and in-vitro cultures of 3T3-L1 adipocytes. Approximately 5-6×10^6^ adipocytes from tissue and 3×10^6^ 3T3-L1 adipocytes were used in this study. The isolated adipocytes were washed with chilled DPBS and lysed using freshly prepared lysis buffer. A total 200 μg protein from the adipocyte lysates were either incubated with 3 μg of HIF-1α/FetA or HIS_6x_/HA antibodies overnight at 4°C using a rotating mixture platform. The immunocomplexes were precipitated using protein-A magnetic beads (surebeads^TM^, BioRad) for 2 hours and washed with 0.1% Tween 20 containing DPBS. This process repeated thrice and the immunocomplexes were resuspended in 4x reducing laemmlli sample buffer and boiled for 5 minutes. The supernatant was collected and subjected to SDS-PAGE followed by immunoblotting analyses with corresponding antibodies.

#### Immunoblotting

Total protein isolation and immunoblotting was performed following previous studies(Choudhary et al, 2023). Briefly, cells were lysed and centrifuged at 13,000 rpm for 10 min at 4°C. Protein concentrations of cell lysates were determined by using the BCA Protein Assay Kit following manufactures’ guidelines. Protein isolation from the adipose tissue was performed by weighing 300 mg of tissue and lysing it in RIPA lysis buffer (for 3 times and 30 seconds each) and shaking in Tissue lyser (Qiagen). Cell lysates (50μg of protein) were resolved on 10% SDS–PAGE and transferred onto PVDF membranes using Trans-Blot Turbo System (Bio-Rad Laboratories). Membranes were first blocked with 5% BSA in TBST (Tris-buffered saline+0.05% Tween 20) buffer for 1h followed by the overnight incubation with primary antibodies in a rotating shaker at 4°C. The membranes were then washed three times with TBST (TBS containing 0.1% Tween 20) wash buffer for 10 min intervals and incubated with peroxidise conjugated specific secondary antibodies for 2 h at room temperature. Membranes were then washed three times with TBST for 10 min intervals and subjected to ECL Substrate (Bio-Rad) incubation for 5 min at room temperature. Protein bands were visualized in Chemidoc XRS+ System (Bio-Rad Laboratories, Hercules, California, USA) using Image Lab Software.

#### PAS Staining of SDS-PAGE Gels

Following electrophoresis of protein samples, gels were immediately fixed in a 12% trichloroacetic acid (TCA) solution at room temperature for 30–60 minutes to immobilize the proteins. After fixation, the gels were transferred to a fresh container containing 1% periodic acid in 1–3% acetic acid and incubated at room temperature for 60–90 minutes to oxidize carbohydrate groups. Gels were washed 2–3 times with 15% acetic acid, each wash lasting 5–10 minutes to remove residual oxidizing agents. The gels were then incubated with Schiff’s reagent at 4°C for 1–2 hours to selectively stain glycoproteins. Finally, excess staining reagent was removed by thorough rinsing with distilled water. The stained gels were visualized using a gel documentation system.

#### Senescence-associated beta-galactosidase (SA-β-gal) assay

SA-β-gal assay was performed following a published protocol(Dimri et al, 1995) with slight modifications. Briefly, control and treated adipocytes were washed twice with DPBS and incubated with a fixative solution containing 0.2% glutaraldehyde and 2% formaldehyde in PBS at room temperature for 5 minutes. Cells were then washed thrice with PBS and incubated with a freshly prepared β-gal solution for 8 h at 37°C in a humidified chamber.

#### Immunofluorescence staining

Control and treated adipocytes were washed with PBS and fixed with ice-cold methanol for 5 min. Cells were then permeabilized with 0.1% Triton X-100 in PBS for 10 min at room temperature and blocked with 1% BSA in PBS containing 0.02% Tween-20 for 30 min at room temperature. The primary antibodies were incubated with the permeabilized cells for 1 h at room temperature, washed thrice with ice-cold PBS for 5 min each, and then incubated with fluorochrome-conjugated secondary antibodies for 1h at room temperature in the dark. Before mounting on a glass slide, cells were washed thrice for 5 min each with ice-cold PBS. Coverslips were mounted onto glass slides using anti-fade mounting medium with DAPI. Details of Antibodies are presented in the materials Table.

#### Adipose tissue Processing and cryosectioning

Visceral adipose tissue samples collected from human subjects and mice models were immediately washed in sterile saline and then placed in Neutral buffer formalin (10%) for overnight fixation at 4°C. After fixation, adipose tissues were passed through increasing concentrations of sucrose solutions (10%, 15%, 20%), embedded in OCT (optimal cutting temperature compound, Sigma), and then frozen at -60°C. Tissue cryosections were prepared by using a Cryo-microtome set to an internal temperature below -25°C (Leica CM 1860, Leica Biosystem, Wetzlar, Germany). Immunostaining was performed on tissue cryosections using specific antibodies.

#### Immunostaining and microscopy

Briefly, tissue cryosections (10-12 μm) were placed in gelatin-coated glass slides, fixed in ice-cold methanol for 5 minutes, blocked with 5% BSA containing blocking buffer, and incubated with specific primary antibodies for 1 h at room temperature. After washing, the signal was visualized by subsequent incubation with appropriate fluorochrome-conjugated appropriate secondary antibodies and counter-stained with anti-fade mounting medium containing DAPI. Images were captured either by an inverted fluorescent microscope (Leica DMi8, Germany), with image analysis was performed using LAS X software, or by a confocal microscope (LSM 880 Carl Zeiss, Germany), with image analysis performed using Zen software.

#### H&E staining and imaging

The visceral adipose tissue from animals was collected and subjected to histopathological analysis. Cryo-sectioning of the tissues was performed, as previously described in this manuscript. These tissue sections were placed on gelatin-coated glass slides and subjected to regressive staining using the following steps: 100% alcohol was passed over the sections for 20 seconds, repeated twice; followed by 90% alcohol for 20 seconds, repeated twice; 80% alcohol for 20 seconds; 70% alcohol for 20 seconds; 50% alcohol for 20 seconds. Subsequently, the slides were rinsed with dH2O for 1 minute. Next, the slides were incubated in Hematoxylin for 3 minutes, followed by a 2-minute water rinse. The slides were then dipped three times in a 0.3% acetic acid-containing alcohol solution, rinsed in dH2O, and further dipped several times in 0.3% ammonium water, resulting in bluing, upon rinsing with water. Afterward, the slides were passed through 80% alcohol for 20 seconds, followed by staining with 2% Eosin for 30 seconds. To remove excess staining, the sections were washed with 95% alcohol for 20 seconds, repeated twice, and then with 100% alcohol for 20 seconds. Finally, the slides underwent a few brief dips in xylene before mounting using DPX solution. Subsequently, the H&E-stained slides were examined using a Leica DMi8 microscope for imaging.

#### RNA extraction and Quantitative PCR

Total RNA was extracted from the adipocytes using TRIzol^TM^ (Invitrogen) following the manufacturer’s instructions. The RNA concentration was measured by using NanoDrop^TM^ OneC spectrophotometer (NanoDrop Technologies, Thermo Scientific, Waltham, MA USA) and 100 ng of RNA was then treated with DNase I and reverse transcribed using the iScript^TM^ cDNA Synthesis Kit (Bio-Rad, Hercules, California, USA). We used PowerUp^TM^ SYBR Green Master Mix (Applied Biosystems) to perform real-time quantitative PCR in QuantStudioTM3 Real-Time PCR System (Applied Biosystems, Waltham, Massachusetts, USA) using gene specific primers. After the final extension, a melting curve analysis was performed to ensure the specificity of the products, and the fold changes in expression were determined using 2^−ΔΔ^CT method. The mRNA expressions were normalized to β-actin. Primer sequences used in this study are listed in Expanded View Table 1.

#### In-silico docking analyses

The proteins (HIF-1α and FetA) structures were sourced from the European Bioinformatics Institute’s AlphaFold Protein Structure Database (EMBL-EBI)(Jumper et al, 2021). These structures were subsequently prepared using PyMol (PyMOL v2.5-Incentive Product, Schrodinger, LLC). Later Then the protein-protein interaction study analysis was performed by ClusPro, a web server utilizing the PIPER rigid-body docking method(Kozakov et al, 2017). This process involves initially sampling all translational and rotational orientations of a ligand-protein concerning a receptor protein. Subsequently, it employs the fast Fourier transform (FFT) correlation approach, utilizing knowledge-based or statistical potentials as the scoring function.

This allows the software to organize the samples and select the most suitable model for the complex. The ClusPro server conducts three computational steps: 1) Rigid-body docking: It samples an extensive number of conformations, numbering in the billions. 2) Root-mean-square deviation (r.m.s.d.)-based clustering: This identifies the largest clusters among the 1000 lowest-energy structures, representing the most probable complex models. 3) Refinement: Selected structures undergo energy minimization. The efficiency of this method lies in its ability to efficiently calculate energy functions using FFTs. This allows it to thoroughly sample countless conformations of the interacting proteins, evaluating energies at each grid point. Consequently, ClusPro can perform protein docking without prior knowledge of the complex’s structure. The obtained results underwent analysis and refinement through PyMol.

#### Luciferase reporter assay

The HIF-1α promoter luciferase assay was performed as previously described(Patra et al, 2023). Briefly, 3T3-L1 mature adipocytes were co-transfected with Firefly HIF-1α promoter plasmid (3 μg/well) and with either AHSG OE plasmid (2 μg/well), or AHSG ASO (100 nM/well) using Lipofectamine LTX with PLUS reagent in 12-well plate for 48 h. Cells were then treated with or without palmitate (0.75 mM) and hypoxia (1% O_2_) for 16 h. On termination of incubations, adipocytes were lysed, and luciferase activity was determined by using Rapid Detection of Firefly Luciferase Activity Kit (E1500, Promega) in Glo-max Navigator Microplate Luminometer (Promega) following to the manufacturer’s protocol. Data was normalized by co-transfecting cells with a Firefly plasmid (2 μg/well) as an internal control. Three independent experiments were quantified for all treatments.

#### Site-directed mutagenesis

AHSG-bio-His plasmid was used as a template for the generation of FetA mutants and HA-HIF1alpha-pcDNA3 was used as template for the generation of HIF-1α mutants using a QuickChange Lightning mutagenesis kit following the manufacturer’s protocol. Primers used to generate the specific FetA and HIF-1α mutants were designed with the help of QuickChange Primer Design Program available at Agilent website (<u>QuickChange Primer Design</u>). Forward and reverse primer sequences used to generate mutation are listed in Expanded View Table 1.

#### Affinity purification for FetA and HIF-1α

HEK-293T cells were transfected with WT and mutant AHSG-bio-His plasmids using Lipofectamine 3000 following the manufacturer’s protocol. 48 h upon transfection cells were harvested in RIPA buffer containing protease and phosphatase inhibitors and subjected to sonication using Sonic Ruptor 250 (omni-international) for thorough lysis. Cell lysate was used for affinity purification of FetA and its mutants by HisPur^TM^ Ni-NTA Resin following manufacturer’s instructions. Briefly, cell lysate (100 ug of protein) was mixed with equal volume of equilibration buffer to prepare the sample. Prepared protein sample was then added to the resin column. His-tagged WT-FetA and its mutant proteins were then eluted using elution buffer from separate resin column setup.

HEK-293T cells were transfected with WT and mutant HA-HIF1alpha-pcDNA3 plasmids using Lipofectamine 3000 following the manufacturer’s protocol. After 48 h of transfection, cells were harvested in RIPA buffer containing protease and phosphatase inhibitors and subjected to sonication using Sonic Ruptor 250 (omni-international) for thorough lysis. The cell lysates (100 ug of protein) were subjected to immunoaffinity chromatography using NHS-Activated Agarose Resin as matrix.

Briefly, NHS-Activated Agarose Resin was coupled with 500 ug of anti-HA antibody for 2 h at room temperature, packed in column then blocked with 1 M Tris, pH 7.4, overnight at 4°C. Unbound antibodies were washed using wash buffer (PBS, pH 7.4). The columns were then incubated with the protein sample overnight at 4°C, followed by two-time PBS wash. HIF-1α and its mutants were eluted using elution buffer (0.15 M glycine HCl, pH 2.5) followed by immediate neutralization of pH by 1 M Tris-Cl, pH 9. Eluted proteins are passed through a desalting column (PD-10) prior to quantification.

#### SPR analysis

The surface plasmon resonance (SPR) analysis was performed to measure the binding affinity between WT/mutant FetA and HIF-1α protein. The measurements were performed using a Biacore-X100 instrument using either NTA or CM5 sensor chips (GE Healthcare). All the measurements were performed using 20 mM HEPES buffer containing 150 mM NaCl, pH 7.3 at 25 °C. The surface of the CM5 sensor chip was coated (flow rate 10 µl/min) with anti-His-antibody as the target proteins contained a His_6x_ tag. The anti-His-antibody capture level was 7000 response units (RU). One of the proteins (His-tag) was then immobilized on the antibody-coated surface as a ligand, and the other protein (without His-tag) was then passed over the surface as an analyte and vice versa. BSA was chosen as a negative control. The sensorgrams were analyzed to investigate the corresponding equilibrium dissociation constant (K_D_) of the protein-protein interaction.

#### Chromatin Immunoprecipitation (ChIP)-PCR

ChIP assay was performed using the SimpleChIP® plus Enzymatic Chromatin IP kit (Magnetic Beads, Cell Signalling technology, 9005). Briefly, upon confluency of 3T3-L1 preadipocytes in 100 mm tissue culture plate, cells were stimulated for differentiation into mature adipocyte, as previously described(Banerjee et al, 2022). The mature adipocytes were then treated with or without palmitate (0.75 mM) and hypoxia (1% O_2_) for 16 h. On termination of incubations, adipocytes cells were processed according to the manufacturer’s protocol. The crosslinked fragmented chromatin was immunoprecipitated by using anti-HIF-1α antibody. The DNA was purified and eluted DNA was then performed for PCR analysis using specific primers flanking the promoter region of *p53* and *Glb1*, then followed by agarose gel electrophoresis(Singh et al, 2019). The gel band intensity was quantified using ImageJ software by considering the amount of immunoprecipitated DNA in each sample represented as relative expression to the total amount of control input chromatin.

#### Statistical analyses

Data are represented as mean ± SD. Student’s *t*-test, two-way ANOVA were used to determine statistical significance, and a p-value at the level of p<0.05 was considered significant. Student’s t-test was used for the comparisons among two groups and two-way ANOVA analyses were considered for calculating significance when comparing multiple groups. All data analyses were performed using GraphPad Prism software (v.8.0; GraphPad Software, Inc., La Jolla, CA).

